# One-step pipetting of barcoded planar microparticles into compact monolayer assembling chip for efficient readout of multiplexed immunoassay

**DOI:** 10.1101/2022.01.03.474850

**Authors:** Sangwook Bae, Daewon Lee, Hunjong Na, Jisung Jang, Sunghoon Kwon

## Abstract

Barcoded planar microparticles have many qualities suitable for developing cost-efficient multiplexed immunoassays. But at the translational research level, there are a number of technical aspects yet remain to be addressed which includes robustness and efficiency of the assay readout process. Assay readout process involves automated barcode identification and signal intensity values from each planar microparticle. For this, each microparticle has to be correctly aligned for correct barcode readout while being, ideally, compactly assembled for maximum microparticle imaging efficiency. To simultaneously achieve such alignment and assembly of microparticles but in a straightforward manner, we designed a microfluidic microparticle assembling chip that only requires a single pipetting step. Our design utilizes capillary flow based guided particle assembly, which allows maximum microparticle-based immunoassay readout efficiency. With the aid of image processing algorithms, we obtained good multiplex immunoassay readout accuracy similar to conventional imaging platforms. Our approach is applicable to both soft elastomer materials (e.g. PDMS) and rigid materials (e.g. polystyrene), the latter of which is frequently used for injection molding based mass production. We anticipate our device could help developing facile and user-friendly platform technologies based on barcoded planar microparticles.

## Introduction

To date, various suspension array-based assay platforms have been developed primarily to increase the multiplexity of assays. Among those, barcoded planar microparticles have many desirable qualities. First is scalability; the potential number of simultaneous measurement (multiplexity) grows proportionally with the number of encodable barcodes which is significantly higher than most non-barcoded particles or assays^1-6^. Second is generality; due to many chemical modifications and particle material available^7^, these particles can be applied to detecting various analytes including oligonucleotides^8-13^, proteins^14-18^ and cells^19-22^. And last is affordability; adaptation of sensitive colorimetric assays^23, 24^ and inexpensive readout tools such as scanners or smartphones^4, 25, 26^ enables development of compact, facile devices. Therefore, one could rationally argue barcoded planar microparticles are good candidates for developing highly multiplexed immunoassays in an efficient manner.

Although many proof-of-concept studies were done, many still require companion technologies to enable translation to commercial products^27^. In one respect, to make such multiplexed immunoassay simple and efficient, two important yet not well-appreciated steps have to be dealt with; i) efficiently aligning the particles and ii) tightly packing these aligned particles into a single layer on a flat surface. This is because, since these microparticles have planar barcodes, accurate and reproducible reading requires particle alignment. Also, tightly packing these aligned particles is necessary to maximize reading throughput.

To address these particle maneuvering problems (aligning and packing), several methods have been invented including magnetic^1^ fluidic flow-guided^28-33^ and shape-guided^22,33^ approaches. One commercialized platform that utilizes microfluidic flow-guided assembly is Mycartis now available for clinical and pharmaceutical applications (patent EP2684605A1). Such fully automated microfluidic device will find many needs in hospital labs and pharmaceutical companies where routine, large scale immunoassay is necessary.

But one could also anticipate need for multiplexed immunoassays from other research labs or point-of-care device developers. For this, further simplifying these particle maneuvering mechanisms to something plain as pipetting would be desirable. We previously developed a method for one-step pipetting-based alignment and assembly of barcoded planar particles^34, 35^. Although this facile method have great potential for multiplexed chemical reaction screening, the packing density had to be sacrificed to prevent well-to-well cross-reaction.

Here, we present a new chip for simultaneous alignment and packing of barcoded planar microparticles. A single particle pipetting of these particles into the chip leads to their automatic alignment and monolayer packaging into the field of view (FOV) of 2x 2.5mm with maximum density. The mechanism doesn’t need any syringe pumps, controllers or other exterior devices. Users can efficiently obtain multiplexed immunoassay results from the packed particles. Unlike spreading particles on plain surfaces, our chip based method doesn’t suffer from particle clumping, overlapping or density fluctuation. We applied our device to fluorescence based multiplexed cytokine immunoassay and obtained equivalent results to traditional planar assay. Although our initial chip design was based on PDMS, we obtained similar particle packing performances with polystyrene (PS) chips, which is potentially better suited for scale up production and commercialization.

## Materials and Methods

### Materials

PDMS, SU8 2015, SU8 2050, isopropyl alcohol (IPA), and SU8 developer were purchased from Microchem. Perfluorooctyltrichlorosilane, tetraethoxyl silane (TEOS), Bovin serum albumin (BSA), (3-aminopropyl)triethoxysilane (APTES) > 98%, Succinic anhydride (SA) > 99%, Polyurethane (PUA), 3-(Trimethoxysilyl)propyl methacrylate (TMSPA), Trimethylolpropane ethoxylate triacrylate (ETPTA), 2-Hydroxy-2-methylpropiophenone (Darocur), and Triethylamine (TEA) > 99.5% were purchased from Sigma Aldrich. Other chemicals and solvents including N-Hydroxysulfosuccinimide Sodium (NHS) salt 97%, Absolute ethanol, NN-Dimethylformamide (DMF), and 1-(3-Dimethylaminopropyl)-3-ethylcarbodiimide hydrochloride (EDC) 98+% were purchased from Alfa Aesar. Capture antibodies, biotin-labelled polyclonal antibodies, and antigens were purchased from either Millipore, eBioscience, or BD Bioscience. Streptavidin-linked phycoerythrin (SA-PE) was purchased from ProZyme. [2-(N-morpholino)ethanesulfonic acid] (MES) buffered saline pack was purchased from Thermo Scientific. Chemicals sensitive to moisture, such as NHS and EDC, was stored in desiccators. Other reagents were stored according to the manufacturer’s recommendation.

### Chip synthesis

For the PDMS chips, we designed FCG type film masks or chrome masks with Autocad and manufactured by Microtech. On 4 inch silicon wafers, for various SU8 pattern heights, we performed spin coating, soft baking, UV exposure, post-exposure baking was done as the manufacturer’s guide. After developing, hard baking was done at 150°C for 10minutes. To reduce the surface energy, we coated the SU8 patterned wafer with perfluorooctyltrichlorosilane by dispensing few microliters of perfluorooctyltrichlorosilane on slide glasses heated to 85°C and placing the wafer over the slideglass for 1 minute. PDMS (Sylgard 184, DowCorning) was mixed with curing agent at 10 : 1 w/w and poured into the SU-8 mold. After baking at 110° C for 30 min,the PDMS pattern was peeled off and attached to a slide glass by O2 plasma treatment (CUTE-MP, FemtoScience).

For PUA chip synthesis, we spread liquid PUA on a glass mold with a ramp with varying angle then UV cured the liquid into a polymer matrix. Then we assembled these triangular PUA blocks on the SU8 pattern of the original particle packing channel. Using this modified 3-dimensional pattern, we fabricated a series of PDMS molds and a final PUA channel that has a channel inlet height gradient.

For PS chip, we outsourced the injection molding to Quantamatrix using our autocad design for the optimized gradient channel design obtained from PUA chip experiments.

### Particle synthesis

We used the previously reported protocol^36^. Briefly, ETPTA was mixed with 10% w/w TMSPA and Darocur was added at 10% w/w of the ETPTA/TMSPA mixture. the ETPTA/TMSPA/Darocur mixture was poured on air plasma treated hydrophilic glass slides. Using masks for contact printing UV lithography using negative resist was performed. TEOS was used for silica coating. Amine groups were coupled to the particle surface using 0.095g/mL APTES solution (in absolute ethanol) and stirred at 25 °C for 2 hours. Bead were washed with absolute ethanol and dried at 110°C for 10 minutes. Terminal amine carboxylation was done by adding carboxylation solution (6mg SA, 8.4uL of 7.2M TEA in 1ml DMF) and stirring at 25°C for 2 hours. Beads were washed with DMF, absolute ethanol, and MES buffered saline. EDC/NHS-cross-linking solution (5mg NHS, 5mg EDC per 1ml MES) was then added and stirred at room temperature for 25minutes and washed with MES. Capture antibodies in MES (14ug/mL) was mixed with bead and stirred (1200rpm) at room temperature for **2 hours**. After antibody cross-linking, the beads were blocked with BSA solution (1% BSA in PBS).

### Immunoassay

Immunoassay protocol was adapted from a previous report (**Jiyun Kim et al**.,). Particles were incubated with antigens (diluted in 0.1% BSA in PBS) and stirred at 4°C for 2 hours. After washing with washing solution (0.1% BSA, 0.02% Tween 20 in PBS), particles were incubated with 2*μ*g/ml biotin-labelled anti-antigen polyclonal antibodies prepared in dilution solution (PBS solution containing 0.1% BSA) at 650rpm, 25 °C for 1 hour and were washed with washing solution. Finally, particles were incubated with SA-PE solution (1 *μ*g/ml SA-PE prepared in dilution solution) for 30 min and were washed rigorously with washing solution before imaging. We imaged the particles with Olympus fluorescence microscope (4× objective lens) with emission wavelength filtered at 575 nm–625 nm at exposure time of 0.2sec. The immunoassay readouts from either the packing channel or 96wellplate wells were decoded using an in-built program from Quantamatrix Inc.

### COMSOL simulation

We used COMSOL Multiphysics 5.4 for our fluidic simulations. We used the ‘particle tracing for fluid flow’ module with no wall slip condition, extra-coarse meshing, and creeping normal inflow with velocity of 3.5mm/sec. Parameter sweeping was used for either i) sweeping the effects of the parameter (e.g. channel height or gradient angle variation) on particle movements or ii) determining the optimal diverging and converging channel design.

## Results

### Channel design optimization

To assess the basic characteristic of planar particle dispersion, we initially tested simple particle dispersion on 96 wellplate wells. The goal was to obtain as many particles per FOV possible while minimizing any unwanted dispersion patterns like particle overlapping and particle tilting. By testing various particle concentration, we observed the trade-off between particle packing density and particle monolayer assembly rate (**Fig 2c**). To overcome this trade-off, we had to come up with a guided assembly method that enables both particle alignment and dense packing. For this we designed a fluidic channel with particle stopping pillars (**Fig 1**.).

**Figure 1.**
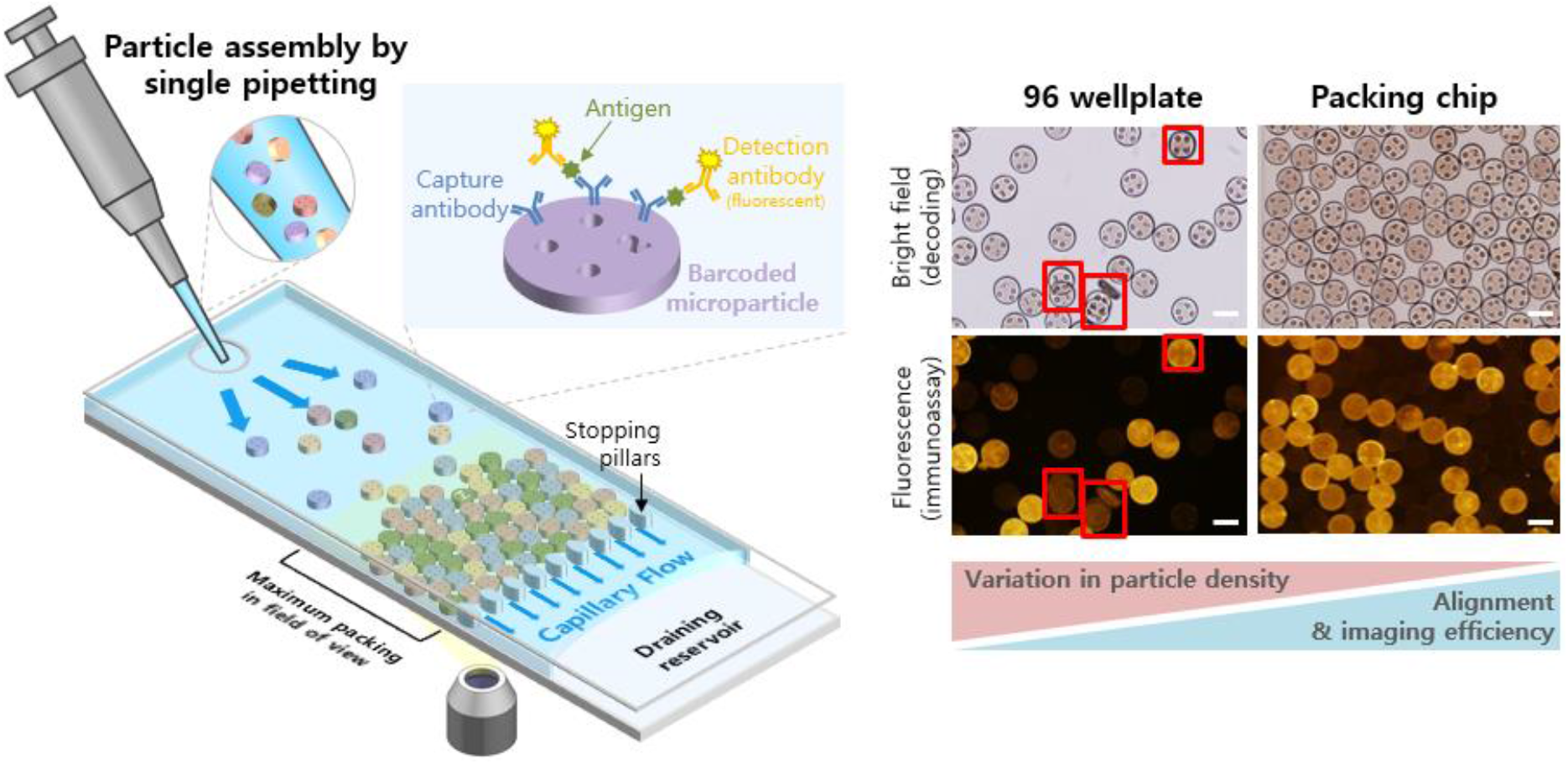
Schematic of a microfluidic, pipetting-based packing of encoded microparticles. Efficient readout of encoded microparticle-based multiplex immunoassay can be done by a pipetting-induced capillary flow-based monolayer assembly of microparticles into a microfluidic chip. The assay readout becomes efficient compared to simple microparticle dispensing on flat surface (i.e. 96 wellplate) because of simultaneous increase of microparticle assembly density and alignment quality. Scale bars are 100μm.

**Figure 2.**
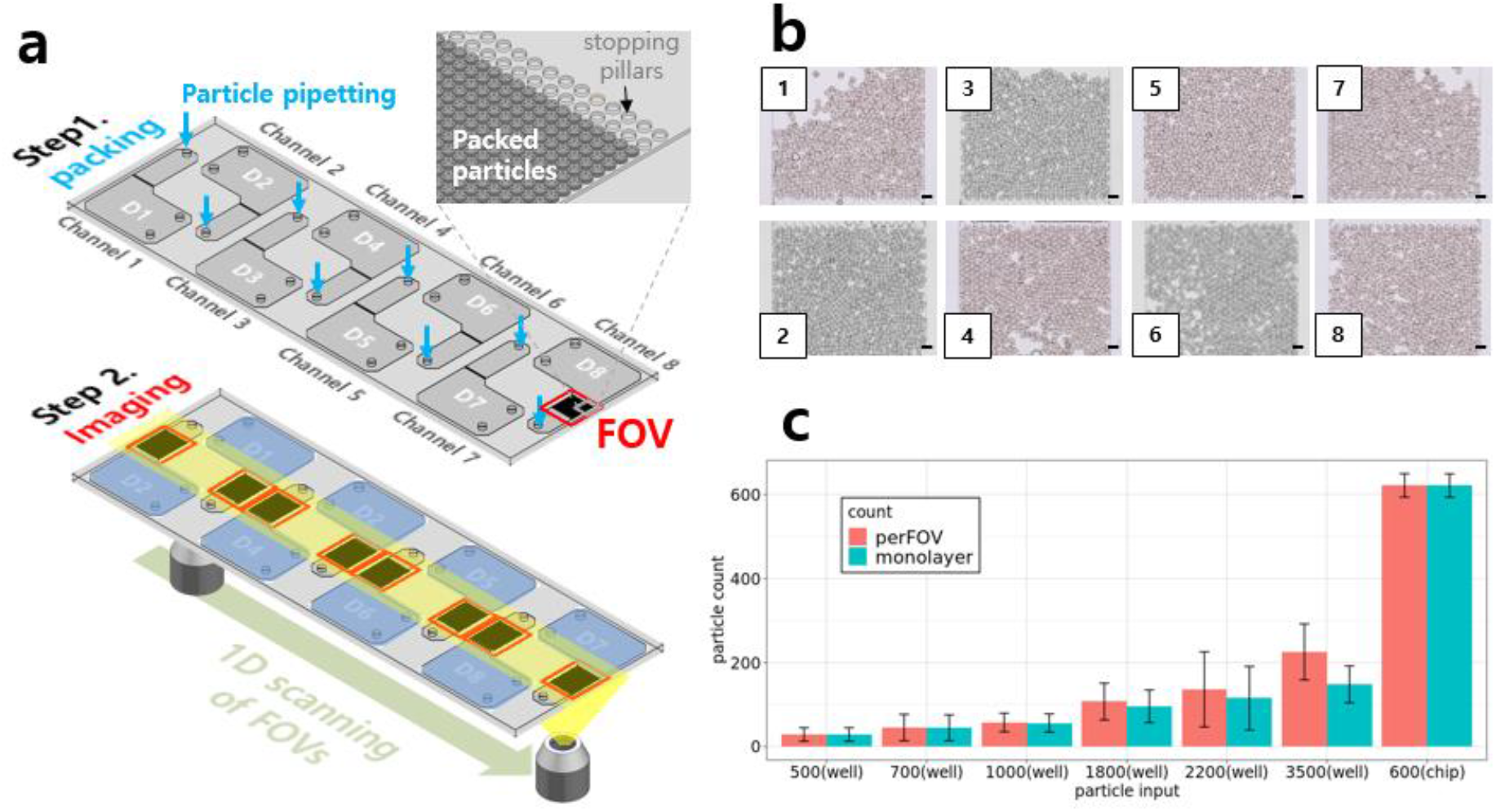
Maximization of imaging throughput using parallel packing channels. (a) Schematic of microparticle assembly and 1D assay readout using an 8 parallel channel chip. After 8 microparticle pipetting into the channel inlets, assay readout can be done in an efficient 1D scanning motion due to the 8 FOVs being aligned in a straight line. (b) Example images of 8 FOVs containing packed microparticles. Scale bars are 200μm. (c) Comparison of particle packing performances between 96 wellplate (well) and particle packing chip. For the 96wellplate experiments, we dispensed either 500, 700, 1000, 1800, 2200, or 3500 microparticles per well. For the packing chip experiment, we pipetted on average 600 microparticles per chip. perFOV (red): number of particles present in a single FOV. Monolayer (blue): number of microparticles per FOV that are assembled into a monolayer without particle overlapping or misaligned. Error bars indicate standard deviation.

With dummy planar particles, we first tested different PDMS channel heights. Our particles were planar cylinders with 100um in diameter and 30um in thickness and were suspended in DIW. Our initial intuition was that channel height should be bigger than the particle height because the suspended particles will have random orientation and thus will need sufficient space to rotate towards proper alignment while moving towards the particle packing region of the channel. However, with increasing channel height, the packed particles will occasionally overlap with adjacent particles which is detrimental to proper signal readout. Our experiment results showed that the channel height has to be nearly identical to the particle thickness (**Fig S1**.) Only then, we were able to get a 99.7% monolayer packing efficiency. The reason these planar particles with 100um diameter were able to enter the 30um height channel was due to the flexible bulging of the PDMS channel (**Fig S2a**). In other words, when we inserted the pipette into the channel inlet and inject the particle-suspended solution, the pressure created a channel height gradient from >100um at the inlet gradually decreasing to 31um at the particle packing region. Harnessing this channel height gradient became the key design factor for our particle packing channel.

With this initial PDMS channel design, we created a chip with 8 parallel channels to increase readout throughput (**Fig 2a**). The channels were arranged in a way that the particle packing regions were aligned in a single line thus enabling easy sequential imaging. We dispensed approximately 600 to 700 particles per channel using normal pipettes. When compared simple particle dispensing in 96wellplates, our PDMS channel was superior in particle packing density as well as monolayer packing efficiency (**Fig 2b, c**). Even with 3500 particle input, 96well results showed 3 times lower number of particles per FOV compared to PDMS channel with 700 particle input. And PDMS channel results showed almost complete monolayer assembly of the particles with only one or two particles per FOV lost during analysis due to particle overlapping. We also compared the particle imaging speed between our channel and 96wellplate dispensing. We designated imaging speed as simply the number of monolayer particles per FOV because the delay between each FOV imaging is comparable between our channel-based method and simple dispensing. Our channel design had one anticipated advantage in speed over simple dispensing due to the straight-line scanning mode rather than the raster scanning of 6∼10 2mm x 2.5mm FOVs per well (d = ∼6.4mm) required to image circular wells. Overall, the image acquisition speed of our channel design was 18∼30 times faster than 96 wellplate due to the significantly reduced number of image acquisition required.

### Multiplexed immunoassay readout

We then analyzed the multiplexed immunoassay readout capability of our channel design. We first created standard curves for both our particle packing channel and 96wells using barcoded particles for detecting IL-1β. We treated our barcoded particles (single barcode) with varying concentration of IL-1 β (0, 800, 4000, 20000, 100000pg/mL) and detection antibodies conjugated with fluorescent dyes. For each IL-1β concentration condition, particles were either injected into our channel or dispensed in 96wells and imaged with a fluorescence microscope. The barcodes and corresponding fluorescence signal of each particle was decoded using a custom software created by Quantamatrix. The standard curves were identical between two particle packing methods (**Fig 3a**). We then performed 4plex immunoassay of cytokines. We prepared 4 types of barcoded particles with each having capture antibodies against IL-2, IL-4, IL-6, IL-5 respectively. We treated these particles with a mixture of cytokines with pre-determined concentrations; 10000pg/mL for IL-2, 2000pg/mL for IL-4, 400pg/mL for IL-5 and 0pg/mL for IL-6. We then treated these particles with detection antibodies conjugated with fluorescent dyes. These particles were then either injected into our particle packing channel or dispensed on 96wellplates. Similar to the singleplex experiment, the assay result was overall comparable between 96wellplate and our particle packing channel (**Fig S3a**). However, our particle packing channel had a slight tendency of reduced signal at high antigen concentration (IL-2 and IL-4) while producing increased signal at low antigen concentration (IL-5 and IL-6). Most critically, our particle packing channel induced a false positive signal for the negative control (IL-6).

**Figure 3.**
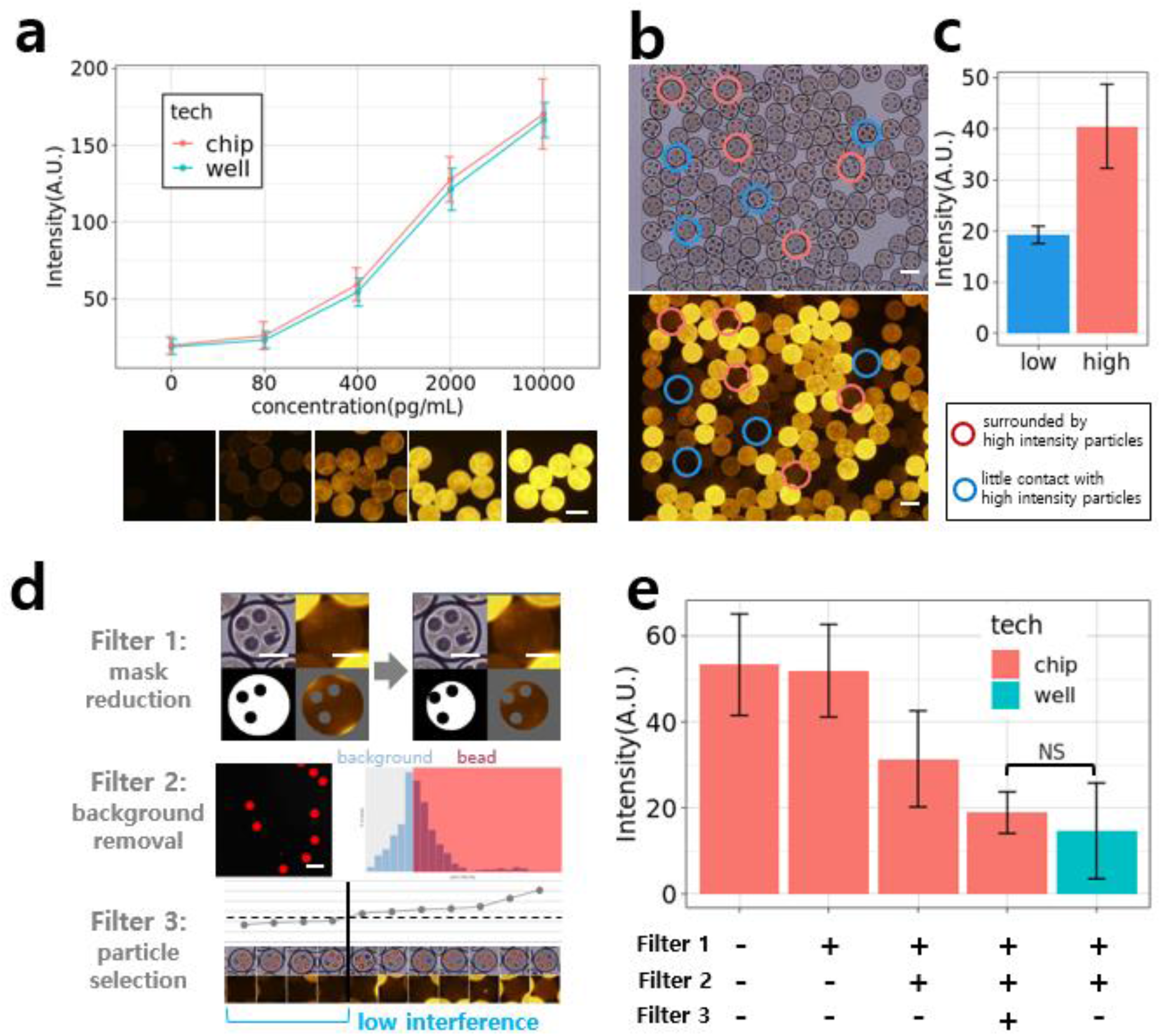
Denoising the fluorescence crossover occurring in packed microparticles. (a) Standard curve of IL-1β immunoassay obtained from imaging singleplex immunoassay microparticles placed on either 96wellplates or in microparticle packing chips. Different code indicated different input IL-1β antigen concentration. Scale bar is 100μm (b) Bright field (upper panel) and fluorescence (lower panel) images of an example multiplex immunoassay readout, where different encoded microparticles showing different fluorescence intensity (or input antigen concentration). Red circles indicate low fluorescence intensity microparticles surrounded by high intensity microparticles. Blue circles indicate low fluorescence intensity microparticles not surrounded by high intensity microparticles. Scale bars are 100μm. (c) Average fluorescence intensities of the microparticles indicated in (b). Error bars are standard deviations. (d) Fluorescence intensity filtering strategy for low fluorescence intensity microparticles surrounded by high intensity microparticles. Scale bars are 50μm for images in filter 1 and 200μm for filter 2. (e) Assessment of filter combinations on fluorescence intensity correction. The microparticle fluorescence intensity values obtained from 96wellplate experiments (but with additional application of filter 1 and 2) was used as a true value reference.

### Correction of fluorescence profile obtained from particle packing channel

We analyzed this discrepancy between 96well and particle packing chip experiments. After screening each particle images decoded by the software, we noticed that when particles with low signal are placed adjacent to high signal particles, signal overestimation occurred in the low signal particle (**Fig 3b, c**). And the magnitude of such signal overestimation correlated with the number of adjacent high signal particles (**Fig S3b**). So we concluded the cause of such signal overestimation was fluorescence crossover between adjacent particles. To remove such errors, we introduced imaging processing techniques for error filtering (**Fig 3d**). We first assessed the effect of reducing the mask size from 100% down to 80% and 60% of the original mask radius. The mask size reduction had a slight effect of reducing the overestimated signals for low signal particles. We then reanalyzed the 96well and particle packing chip experiment images using these modified mask sizes. Although we reduced the signal overestimation for low signal particles in the packing channel, it was still higher than the 96well results. Then, to remove the image background signals, we masked the particles, measured the background pixel intensities, removed pixels with intensity above the median value, calculated the average of the remaining pixels, and subtracted this average value from all particle intensity values. We also assessed the effect of guided subsampling of particles. This was introduced to remove particles that have adjacent particles with high signal particles. Due to limited sample size, this subsampling method was done heuristically. But future large collection of images will help establish more objective and statistically defined thresholds for this process. We found that by applying all three filtering procedures above, the assay profiles between packing chip and 96wells becomes comparable. Most importantly, signal overestimation of the low signal particles in packing chips was successfully removed (**fig 3e**). By applying the three filtering steps above to our previous 4plex immunoassay result, we were able to obtain similar assay profile between our packing chip and 96well experiments (**Fig 4c, d**). It should be noted that for the IL-1β standard curve generation experiments, unlike singleplex experiments in Figure 3a requiring 5 independent experiments required, the standard curve in Figure 4c was obtained by a single experiment using 5plex assay.

**Figure 4.**
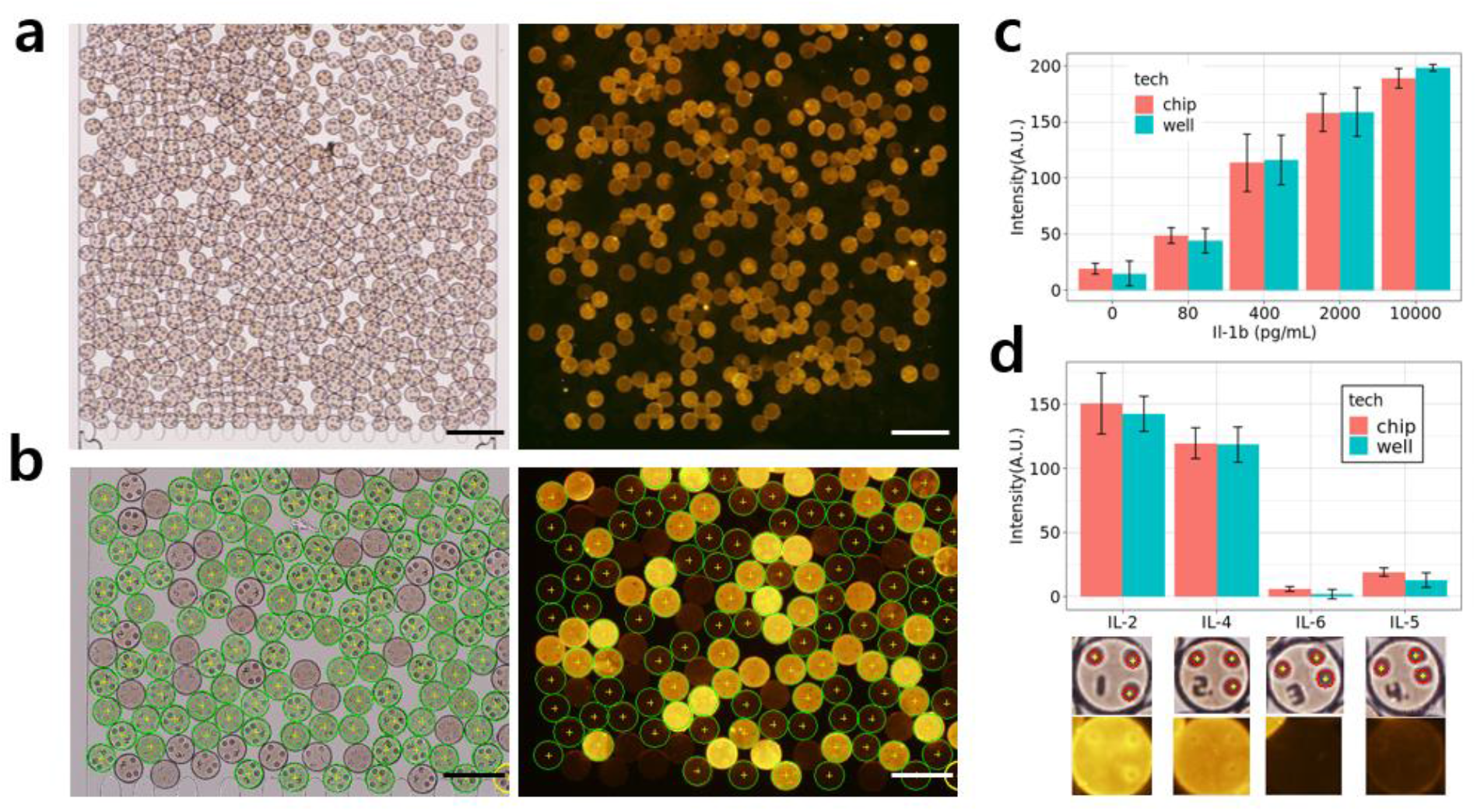
High throughput multiplexed immunoassay using packing chip. (a, b) Bright field (left panel) and fluorescence (right panel) images of a 4plex cytokine immunoassay using microparticle packing chips. As indicated in (b), each microparticles were correctly captured by our image processing algorithm. Scale bars are 400μm (a) or 200μm (b). (c,d) Comparison of 5plex one-pot generated standard curve of IL-1β (c) and 4plex cytokine detection profiles (d) obtained by either 96wellplate experiment (blue) or microparticle packing chip (red). Input antigen concentrations were 10000pg/mL for IL-2 (code 1), 2000pg/mL for IL-4 (code 2), 400pg/mL for IL-5 (code 4) and 0pg/mL for IL-6 (code 3). Error bars are standard deviations.

### Particle packing chip design with PUA and PS

We then decided to design a particle packing chip that would be suitable for industrial manufacturing such as injection molding. As mentioned earlier, the channel height gradient design had to be applied. To determine the effective channel height gradient design, we tested a number of approaches. First we tested PUA-based channels having stepwise channel height reduction. Although our fluidic simulation predicted no significant turbulence (**Fig S4a**), the particle packing experiments with this design ended up with particle clogging at the channel height varying junctions (**Fig S2b, c**). COMSOL simulation showed that replacing the stepwise channel height reduction with a continuous height reduction (i.e. gradient channel height) could reduce the particle clogging problem (**Fig 5a, b**). So we tested gradient channel height designs with different gradient angles from 0.5° ∼ 10°. We empirically determined that the gradient angle have to be smaller than 2.5° to effectively prevent particle clogging (**Fig S2b, c**). We also found that the width of the channel have to be wider at the end of the channel height gradient to prevent particle clogging (**data not shown**). To obtain further intuitions for the optimum channel design, we performed COMSOL fluidic simulation with parameter sweeping setup for the width of the end of the gradient channel. Using the particle tracing module we found that a certain diverging-converging channel width structure was needed to prevent particle clogging while minimize dead volume (**Fig 5c, Fig S4b**). In other words, sufficient divergence was needed to prevent particle clogging (or particle sticking to channel wall) while excess divergence leads to dead channel volume where no particles pass through. Based on this optimized design, we then created a mold for PS injection molding. The resulting particle packing PS chips were tested with fluorescent barcoded particles and were successfully imaged (**Fig 5d, e**). The barcodes were clearly readable by human eye even though the transparency of the PS chips were poor compared to PDMS channels. However, the particle decoding programs were not compatible with these images due to the frequent defects on the surface of these PS chips. Future optimization of these injection molding method will be required to reduce these channel defects.

**Figure 5.**
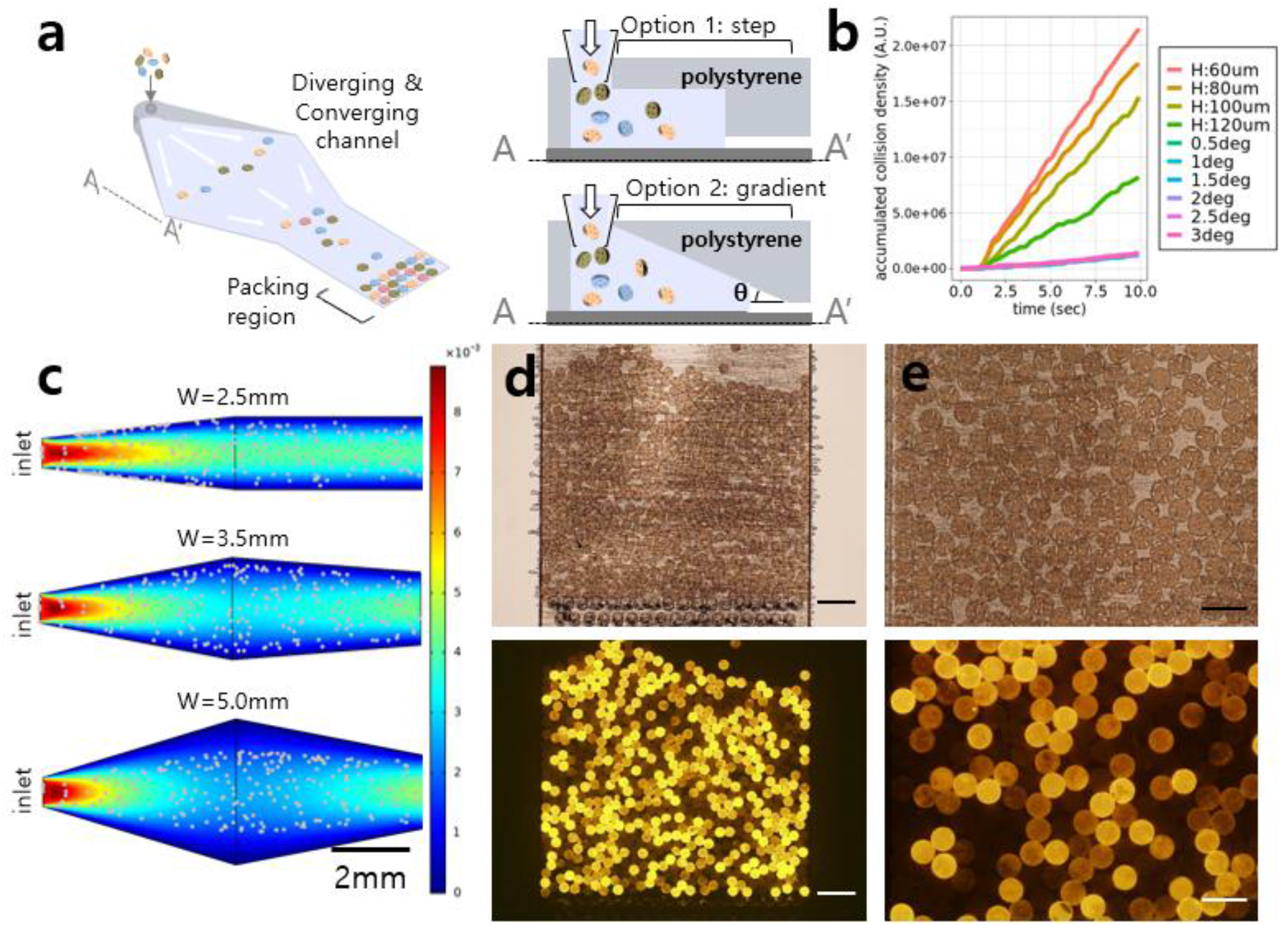
Geometrical optimization of commercializable high stiffness material based fluidic channel. (a) Design of a solid, injection-molded, polystyrene, microparticle packing chip. The diverging and converging channel width and continuously reducing channel height was the key features to enable microparticle packing in such solid-sate microfluidic chips. (b) Simulation result comparing the microparticle channel adhesion density for various channel designs; the channel height profile of these designs were either a step-wise reduction or a continuous reduction with a certain gradient angel. Microparticle channel adhesion density indicates the number of microparticles adhere to the channel wall per unit time per unit area; a parameter we used to assess the microparticles’ channel clogging effect, which must be minimized for efficient microparticle assembly. (c) Simulation result comparing channels with different channel width profiles on the microparticle dispersion efficiency. Color scheme indicates fluidic flow velocity. Each dot represents a microparticle. W indicates the maximum channel width. (d, e) Bright field (upper panels) and fluorescence (lower panels) images of multiplex immunoassay microparticles assembled in a PS microparticle packing chip. Scale bars are 400μm (d) and 200μm (e).

## Conclusion

We created a particle packing chip that is capable of simultaneous particle alignment and monolayer packing of non-spherical, planar microparticles. Performing barcoded microparticle immunoassay with our packing chip, compared to conventional wellplate, leads to 18∼30 times faster assay readout with minimal data loss occurring from improper particle orientation. Other than the chip and a pipette, our strategy doesn’t require cumbersome syringe pumps or other fluidic controlling modules. We expect this chip design will find utility in point-of-care applications where tools are encouraged to be simple yet high throughput, multiplexing capacity is required.

